# Trade-offs in a reef-building coral after six years of thermal acclimation

**DOI:** 10.1101/2023.07.20.549699

**Authors:** Anna Roik, Marlene Wall, Melina Dobelmann, Samuel Nietzer, David Brefeld, Anna Fiesinger, Miriam Reverter, Peter J. Schupp, Matthew Jackson, Marie Rutsch, Julia Strahl

**Author notes:** equal contribution (Anna Roik, Marlene Wall, Julia Strahl).

## Abstract

Evidence is growing that reef-building corals have the capacity to acclimate to new and challenging thermal conditions by increasing their thermal resistance. This raises hopes for their future persistence in a warming world. However, potential trade-offs that accompany such resistance gains, have remained largely unexplored. We provide the first report on the physiological trade-offs in a globally abundant and ecologically relevant coral species (*Pocillopora acuta)*, after a long-term exposure to an elevated temperature of 31 °C in comparison to conspecifics cultivated under a cooler ‘control’ thermal regime. At both temperatures, corals consistently appeared to be visually healthy throughout a six-year period. At 31 °C, corals had increased metabolic rates (both respiration and photosynthesis) that resulted in higher biomass accumulation and total energy reserves compared to the corals from the ambient regime. Further, the composition of coral host tissues shifted in favor of lipid build-up, suggesting an altered mechanism of energy storage. The increase in biomass growth came at the cost of declining skeletal growth rates and the formation of higher density skeletons. In the long-term, this trade-off will result in lower extension rates that can entail major ramifications for future reef building processes and reef community composition. Moreover, symbionts at 31 °C were physiologically more compromised with overall lower energy reserves, possibly indicating a stronger exploitation by the host and potentially a lower stress resilience. Our study provides first insights into a successful thermal acclimation mechanism that involved the prioritization of energy storage over skeletal growth, entailing higher demands on the symbionts. Our observation in this 6-year study does not align with observations of short-term studies, where elevated temperatures caused a depletion of tissue lipids in corals, which highlights the importance of studying acclimation of organisms over their relevant biological scales. Further investigations into trade-offs at biologically relevant scales and how they unfold under an acute heat stress will help to provide a more comprehensive picture of the future coral reef trajectory. Importantly, these insights will also help improve interventions aimed at increasing the thermal resilience of corals which anticipate to use thermal preconditioning treatments for stress-hardening.

**Graphical abstract:** 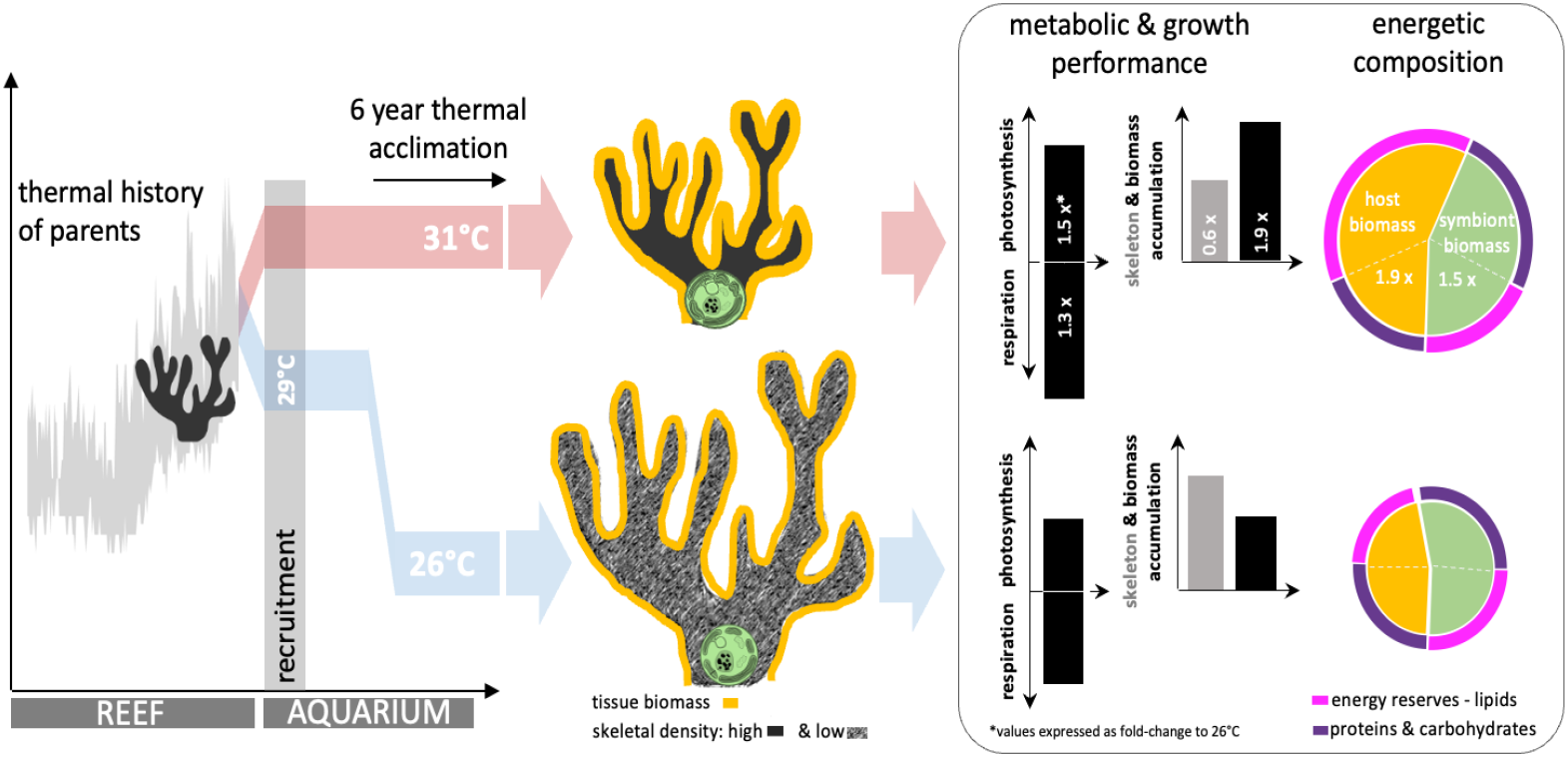

## Introduction

Tropical coral reef ecosystems are facing a major crisis predominantly caused by rising ocean temperatures that lead to coral bleaching, mortality, and reef habitat erosion (Berkelmans et al., 2004; Fukunaga et al., 2022; Heron et al., 2016). The decline of reef ecosystems does not only lead to the loss of reef-building corals, but also of numerous reef-associated marine species. Additionally, serious consequences arise for coastal communities and nations that depend on reef ecosystem services, in particular coastal protection, marine foods, and tourist economies (Eddy et al., 2021; IPBES, 2019). Concerns about the future of coral reef ecosystems have fueled the quest for solutions. Most notably, the umbrella-term “assisted evolution” comprises several innovative ideas of human interventions that aim to help accelerate adaptation and acclimation of reef-building corals to sustain tropical reef ecosystems under future climate change scenarios (van Oppen et al., 2015; Voolstra et al., 2021). Among others, two key strategies are on the rise, one of which builds on (evolutionary) adaptive mechanisms of marine organisms (Elder et al., 2022; Humanes et al., 2021; Kenkel & Matz, 2016) and the other relies on their physiological plasticity and acclimation potential (DeMerlis et al., 2022; Henley et al., 2022; Majerova et al., 2021; Martell, 2023). While many findings indicate that thermal tolerance of corals can be partially explained by genetic variation and, hence, is ingrained in genomes and heritable traits (Howells et al., 2022), some of the unexplained variation in thermal tolerance could be attributed to plasticity (Kenkel et al., 2015; Thomas et al., 2018). It also became obvious that not only the genotype but in large parts environmental impulses drive plasticity (Barshis et al., 2010). To harness coral plasticity, and thus coral acclimation potential, “preconditioning” treatments that expose coral propagules to stressors (or sub-optimal/challenging conditions) have been proposed. This approach aims to prime the corals for stress resistance and has inspired many experimental studies in recent years (Bellantuono, Granados-Cifuentes, et al., 2012; DeMerlis et al., 2022; Henley et al., 2022).

While adaptation through trait selection is a lengthy process that requires generations of organisms to act on, some corals have indeed demonstrated a higher stress resistance compared to others, as well as the capacity to enhance this resistance within a lifetime. This phenomenon has been mostly observed in corals with a history of challenging thermal exposures or experience of highly variable environmental conditions in intertidal reefs, lagoonal reefs, or areas exposed to frequent upwelling (Brown et al., 2002; Buerger et al., 2015; Castillo et al., 2012; Oliver & Palumbi, 2011; M. Wall et al., 2023). Studies have increasingly corroborated that corals, pre-exposed to challenging conditioning and stressors, are likely to perform better under new events of stress compared to those without such pre-exposure, indicating that plasticity (in particular the thermal tolerance range) of corals can be expanded through “environmental priming” (Hackerott et al., 2021; Martell, 2023). Furthermore, data collected throughout temporal (or seasonal) stress events, such as moderate heat waves, have shown that coral survivors were increasingly associated with even higher stress resistance following such events (Ainsworth et al., 2016; Bellantuono, Hoegh-Guldberg, et al., 2012; M. D. Fox et al., 2021). Therefore, physiological acclimation capacity within the lifetime of organisms should be considered as an increasingly important survival strategy for coral species under the environmental changes expected in the coming years and decades.

The prospects for employing thermal preconditioning treatments to generate thermally acclimated corals are promising, but so far it remains poorly understood how trade-offs are associated with gains in thermal stress resistance. Higher temperatures pose physiological challenges for organisms, raising biochemical reaction rates and at the same time increasing energetic demands (Angilletta et al., 2004; Hornstein et al., 2018). Organisms often shift their metabolic strategies as a compensatory response under new thermal conditions, which entails changes in metabolic enzyme activity, modifications in tissue biochemistry and ultimately resource allocation (Tattersall et al., 2012). There is evidence of such metabolic shifts in corals exposed to high temperatures. For instance, Gibbin et al. (2018) have shown how carbon and nitrogen uptake of symbiotic dinoflagellates and coral cells has been altered under elevated temperatures, while corals have remained visually healthy - hence, have likely successfully acclimated to the new thermal condition. However, such shifts in metabolic strategy can entail trade-offs. A trade-off by definition is the outcome of the prioritization of one trait or function at the cost of another (Pörtner et al., 2006). Most commonly this relates to the allocation of resources into a specific trait, which, at a specific moment, maintains optimal performance or is important for stress mitigation (Lesser, 2013). For instance, thermal resistance in marine species is often provided at the expense of growth or reproduction, as the energy investments shift towards cell protection and tissue maintenance under stress (Sokolova et al., 2012). Trade-offs of high temperature resistance have been studied and discussed in numerous species (Fusi et al., 2016; Karl et al., 2013; Petes et al., 2008; Roze et al., 2013; Seebacher et al., 2015; Trip et al., 2014), but are mostly understudied in corals. To date, it has been shown that adaptive (and heritable) thermal resistance can be accompanied by trade-offs, such as declines of coral growth rates and tissue lipid content (Bay & Palumbi, 2017; Howells et al., 2013; Kenkel et al., 2015). Another noteworthy finding is that corals with a higher bleaching resistance naturally tended to host lower numbers of symbiont cells in their tissues (Cornwell et al., 2021). The lower symbiont load came at the cost of a decreased growth rate, likely a consequence of a lower photosynthetic output. In contrast, corals in a short-term (5 weeks) marine heatwave experiment did not show any apparent trade-offs regarding fecundity or growth associated with their heat tolerance (Lachs et al., 2023). However, it is uncertain what the consequences of the changes in metabolic strategies will be when corals endure high temperatures over longer periods of months or years.

To shed more light on potential trade-offs of successful acclimation to warmer conditions, we investigated corals over biologically relevant, year-long timescales. Corals were raised and maintained under two thermal regimes in the lab and remained there for six years (31 °C vs. 26 °C). Their parental colonies originated from a thermal regime of ∼29 °C on average throughout the year, experiencing lower daily winter averages of 26 °C and diel fluctuations between 25 - 33 °C across the year. To answer the question whether trade-offs were inflicted with the acclimation process to the elevated temperature regime of 31 °C, we investigated the metabolic performance of host and symbionts, their tissue compositions (i.e, proxy for energetic condition and strategy), as well as tissue and skeletal growth rates (i.e., proxy for ecological success). We aimed to evaluate whether corals that acclimated to 31 °C underwent any metabolic shift or any potential trade-off compared to those acclimated to the cooler temperature regime.

## Materials and Methods

### History of corals

In July 2015, six *Pocillopora sp.*-type colonies were collected at a depth of 1-2 m from Luminao Reef on Guam, USA, (13°27’55.25”N, 144°38’48.84”E). Luminao is a fringing reef which features an annual average temperature of ∼29 °C and experiences midday temperature peaks exceeding ∼31 °C during the hottest month of the year (Supplementary Figure S1). Larvae were released after the new moon in August 2015. Prior to the anticipated night of larval release, mother colonies were each placed into 15 L containers supplied with aeration and with chips of the crustose coralline red alga *Hydrolithon reinboldii* (also collected from Luminao reef). Immediately after the release, most of the larvae settled on the provided settlement chips. The settled polyps were then glued onto plastic buttons on PVC crates and recruits of all mother colonies were mixed. The coral offspring were subsequently transferred to two temperature regimes, ambient (29 °C) and elevated (30 - 31 °C). The setup consisted of 12 flow-through tanks, each holding 69 settled recruits, either maintained at ambient or elevated temperature. In November 2015, recruits were transported to the tropical seawater facilities at the Institute for Chemistry and Biology of the Marine Environment (ICBM) Terramare in Wilhelmshaven, Germany, where they were kept at their respective temperature (29 °C and 31 °C) until August 2016. Recruits maintained at ambient conditions had a higher survival probability than those under elevated temperature. Approximately half a year after settlement, survival of recruits living at elevated temperature had dropped below 50 %, while survival of recruits living at ambient temperature was above that (i.e., ∼60 %). After one year, less than 25 % of the recruits had survived, with significantly lower survival in the group living at elevated temperature (Figure S2). Survival monitoring was halted in August 2016 and corals remained in the same tanks for the following five years, now at a cooler ambient temperature of 26 °C, i.e., corresponding to the lower daily average temperature of their home reef during winter, and at an elevated temperature of 31 °C, i.e., corresponding to the peak daily average temperature during summer (Table 1). The next assessment of coral performance took place in July and August 2021. Until this time point, all offspring colonies were visually healthy. In 2021, the fully-grown adult offspring colonies were identified as the species *Pocillopora acuta* (see details of species identification in Text S1 and Figure S3).

**Table 1.** Physicochemical conditions in the aquaria for long-term maintenance (Aqu.) and the experimental tanks (Exp.).

|  | Temperature<br>[°C] | Salinity<br>[PSU] | Alkalinity<br>[° dKH] | Phosphate<br>[mg/L] | Nitrate<br>[mg/L] |
| --- | --- | --- | --- | --- | --- |
| **Aqu. Tanks**<br>**2016-2021** | ~26 \| ~31 | ~34 | 7-7.5 | 0.1-0.15 | 1-4 |
| **Exp. Tanks**<br>**2021** | 25.42 ± 0.42 \|<br>30.47 ± 0.54 | 33.73 ± 0.25 | 7.72 ± 0.38 | 0.13 ± 0.04 | 1.31 ± 0.95 |

### Set-up for physiological diagnostics

To prepare corals for assessment, fragments of 12 adult *P. acuta* colonies from each temperature regime were distributed across four experimental tanks per ambient (26 °C) and elevated temperature (31 °C), respectively. Six individual fragments of each colony were cemented into “plugs” using aquarium cement (Stone Fix, Aqua Forest, Brzesko, Poland) and a silicone plug mold. In total, 144 fragments were transferred for 54 days onto racks in six experimental tanks (24 fragments per tank) at either 26 °C or 31 °C (72 fragments/treatment, 3 replicate tanks per temperature). The experiment was run with artificial seawater (Tropic Marin® Pro-Reef, Wartenberg, Germany). An automated dosing system supplied calcium, carbonates, magnesium as well as the relevant trace elements via Balling method. Additionally, nutrients such as phosphates and ammonium were added. Dosing was programmed according to the measured concentrations in order to keep water composition stable. Salinity, temperature, nutrient levels (nitrate, phosphate) and alkalinity were measured regularly throughout the experiment (Table 1). The oxidation reduction potential and pH of the system were monitored with a GHL Profilux 3 computer. Tanks were equipped with LED lights (Radion XR15 G4Pro, max. 90 W, EcoTech Marine, USA) and light levels were adjusted to deliver between 130-150 µmol PPFD throughout the day. A flow pump (Turbelle 6025, Tunze GmbH, Germany) provided sufficient water circulation. Water temperatures were maintained with temperature controllers (BioTherm Pro, Hobby GmbH, Germany) and 300 W titanium heaters (Schemel & Goetz GmbH & Co KG, Offenbach, Germany). During a two-month fragment-acclimation phase, temperature was recorded hourly using HOBO Tidbit v2 temperature loggers (Onset, USA), while light intensity and fragment health (algal overgrowth and tissue paling) were measured weekly. To ensure equal light intensity and water movement for all fragments, coral racks were rotated once a week. Corals were fed twice a week with 50 ml of a feeding solution based on clam, squid, fish and phytoplankton concentrate (Tropic Marin ® Phytonic, Wartenberg, Germany). Prior to the live physiological and biochemical assessments coral fragments were not fed for three days.

### Live physiological measurements

To determine metabolic rates (photosynthesis and respiration), one fragment per colony was incubated under controlled conditions under light and dark conditions following the procedure outlined in Strahl et al. (2015). Briefly, coral fragments were incubated in custom-made, clear acrylic incubation chambers (0.21 L). The fragments were mounted into the chamber lid submerged in the respective experimental tank and kept in place by fixation with plastic screws. The incubation chambers were closed underwater and then transferred to a temperature-controlled water bath, either at 26 °C or 31 °C, on top of a magnetic stirring plate (Multi-point Magnetic Stirrer MS-MP8, Witeg, Germany), to ensure continuous water movement to minimize boundary layer thickness. Light incubations were run for ∼ 60 min at midday with a light intensity of ∼ 160 µE. Dark incubations were performed subsequently by acclimating corals to dark conditions for 45 minutes and performing the dark incubation for ∼ 120 min to determine dark respiration rates. Each chamber was equipped with an oxygen sensor spot (PreSens Precision Sensing GmbH, Germany), connected to a multi-channel fiber optic oxygen transmitter system (Oxy 4-Mini, PreSens Precision Sensing GmbH, Germany) and the associated software OXY4v2_30 (PreSens Precision Sensing GmbH, Germany), recording oxygen saturation in mg L^-1^ every 15 seconds. Two multi-channel systems were available to simultaneously incubate a total of six coral fragments and two ‘blank’ chambers (no coral fragment added). The latter are necessary to correct coral metabolic rates for background photosynthesis and respiratory activity of microorganisms in seawater and/or algae growing on the plug. Thus, a total of four incubation runs (including both dark and light incubation) were necessary to incubate all fragments of both temperature regimes by performing one incubation run a day over four days by alternating the temperature treatments. Oxygen sensors were calibrated with O_2_-free (0 % O_2_, sodium dithionite) and air-saturated (100 % O_2_) seawater prior to the four incubation runs. After the dark incubation, the incubated coral fragments were immediately snap-frozen in liquid nitrogen for further biochemical analysis and surface area determination.

The obtained raw data for both net photosynthesis and dark respiration were derived in mg L^-1^ by taking the temperature and salinity during the incubations into account. The respective rates were calculated by linear regression of the oxygen changes using the software R (R Core Team, 2021) and a customized script including a function from the R package rMR v1.1.0 (Moulton, 2018). The slope of the metabolic rates was analyzed for the entire incubation interval except for the first 15 minutes that were excluded from the calculations. Subsequently, the metabolic rates were corrected for the average “background” rates measured by the two blank incubations, as well as for the incubation volume. Finally, rates of net photosynthesis and dark respiration (mg O_2_ cm^-2^ h^-1^) were normalized to the tissue-covered surface area of each coral fragment.

### Biochemical analysis of tissues

To determine the biochemical composition of both coral host and symbiont, coral fragments were processed following established protocols with slight modifications (Bove, 2021; Buerger et al., 2015). Coral tissue was removed from the skeleton using an air gun and filtered seawater. The tissue slurry of each sample was topped up to a total volume of 20 ml and homogenized for 30 seconds with an Ultra Turrax (IKA, USA). Host tissue and symbiont cells were then separated by centrifugation for 10 min at 4,400 rpm and −1 °C (Eppendorf Centrifuge 5702, Germany). Subsequently, the host supernatant (avoiding the algal endosymbiont pellet) was removed carefully and aliquoted for downstream analyses. The symbiont pellet was washed and resuspended in 3.5 ml filtered seawater and similarly aliquoted for downstream analyses (approx. 1-6 ml for biomass determination, 0.5 ml for both protein and carbohydrate analyses and the rest for lipid content).

For biomass determination, each fraction was filtered, using a pre-combusted filter (4 hours at 500 °C, Whatman GF/C, GF Healthcare Life Sciences, United Kingdom) and then dried for 24 hours at 60 °C and weighted, using an electronic fine balance (Sartorius M2P, Sartorius AG, Germany; precision: 0.001 mg). Biomass (= the weight minus the filter) was calculated per surface of the coral in mg cm^-2^.

Protein, carbohydrate and lipid contents were determined next. For protein analysis, a subsample of the respective tissue aliquot (0.025 ml) was used to measure the protein concentration (Lowry et al., 1951). For this, a protein assay kit (DC Protein Assay Kit, Bio-Rad Laboratories Inc., Hercules, USA) and a bovine serum albumin (BSA) standard were used. Measurements were performed using a photometer (UV-1800 spectrophotometer, Shimadzu Corporation, Kyoto, Japan) at 750 nm. To determine carbohydrate concentration, a subsample of the respective tissue aliquot (0.1 ml) was analyzed using the phenol-sulfuric acid methods after (DuBois et al., 1956) with some slight modifications for measurements with a microplate reader and a D-glucose standard. The absorbances of the samples and standards were measured in triplicates at 485 nm on a microplate reader (TriStar LB941 Multimode Reader, Berthold Technologies). The remaining tissue slurries (a minimum of 1800 µL) were used to determine the total lipid concentration in triplicates (600 µL each) using the colorimetric sulfo-phospho-vanillin (SPV) method for microplate measurements after Cheng et al. (2011) with some slight modifications and a corn oil standard (B Bove & Baumann, 2021). The absorbance was measured at 530 nm with the same microplate reader.

Finally, the protein, carbohydrate and total lipid concentrations were converted to kilojoules (kJ; protein: 23.9 kJ/g, carbohydrate: 17.5 kJ/g, lipids: 39.5 kJ/g) after Gnaiger & Bitterlich (1984) and energy reserve concentration (both in mg and kJ) were standardized to the tissue-covered surface area of the corals. The airbrushed coral skeletons were dried overnight in the oven at 60 °C and the surface area was determined using the single wax dipping technique (Veal et al., 2010).

### Measurements of skeletal traits

Growth rates were determined by measuring 1-2 coral fragments per colony following the buoyant weight technique (Jokiel & Guinther, 1978). Briefly, fragments were weighed while submerged in seawater using a microbalance with an underfloor weighing system (Sartorius, BP 210S). Measurements were taken at the corals’ respective thermal condition. Temperature and salinity were recorded for each measurement. Fragments were weighed with and without their “plug” at the start and the end of the experiment (spanning 42 days). Three “empty” plugs per treatment group were measured alongside the coral measurements to account for accretion of the cement “plug” structure in the final calculation of accretion rates. The obtained buoyant weights were converted into skeletal dry weights using the respective seawater density values (calculated from measured temperature and salinity) and an aragonite density of 2.93 g cm^-3^ (Spencer Davies, 1990). The change in dry weight between the start and the end of the experiment was calculated and subsequently divided by the number of experimental days to calculate diurnal accretion rate. Further, values were normalized by surface area of each coral fragment (mg d^-1^ cm^-2^). The surface area values were determined for each fragment using wax dipping technique in a single dip (Veal et al., 2010). Skeletal densities were determined from an additional fragment per colony using the water displacement volume and the weight of the coral skeleton (Strahl et al., 2015). Fragments were soaked in bleach until the coral tissue was completely detached from the skeletons and were dried overnight in the oven at 60 °C. The pre-weighed branches coated with a layer of paraffin wax were submerged in a beaker filled with freshwater and 0.0048 g L^-1^ benzalkonium chloride, which had been added to break the surface tension of the water. To determine the water volume displaced by the coral branches, overflowing water was collected and measured in pre-weighed petri dishes. The accuracy of the method was evaluated by determining the water displacement of plastic cylinders with known volumes ranging from 0.86 cm^3^ to 5.82 cm^3^, with variations between measurements of <5 %. Subsequently, changes in the extension rate (cm yr^.1^) between treatments were assessed and calculated by dividing the net calcification rates determined by buoyant weight and normalized to surface area (mg cm^-2^ yr^-1^) by the skeletal densities (mg cm^-3^). Note this procedure assumes similar calcification rates along the entire surface area of individual fragments. However, in branching species it is known that tips grow faster (up to 13.2 times faster) than the base (Rinkevich & Loya 1984). Thus, the obtained extension rates do not represent absolute rates of the branch tip, but rather provide an estimate how much extension rates will differ between to thermal treatment groups.

### Statistical analyses

Statistical analyses were performed using R (version 4.1.1). Shapiro-Wilk normality tests were used to test for normality and Levene’s test to test the assumption of equal variance of data among two thermal treatments. Where the data met the assumption for parametric tests, t-tests were performed to determine the differences between the two thermal treatments. Non-parametric Wilcoxon-tests were performed to test the treatment-related differences, where the data did not meet the conditions for a parametric test.

## Results

### Metabolic performance

Net photosynthetic rate and dark respiration rate per coral surface area (indicating the overall performance of the holobiont) were significantly higher by 1.5- and 1.3-fold at the elevated temperature, respectively (both comparisons: *p* < 0.05). Both metabolic rates per biomass weight (indicating the performance per tissue unit/ cell unit), however, did not differ between the two temperatures (Figure 1 A & B). The net photosynthetic rate had medians of 0.014 and 0.016 mg O_2_ mg^-1^ biomass and respiration rates 0.005 and 0.004 mg O_2_ mg^-1^ biomass, for 31 °C and 26 °C, respectively. Further, fragments at 31 °C overall exhibited a slightly higher variability.

**Figure 1.**
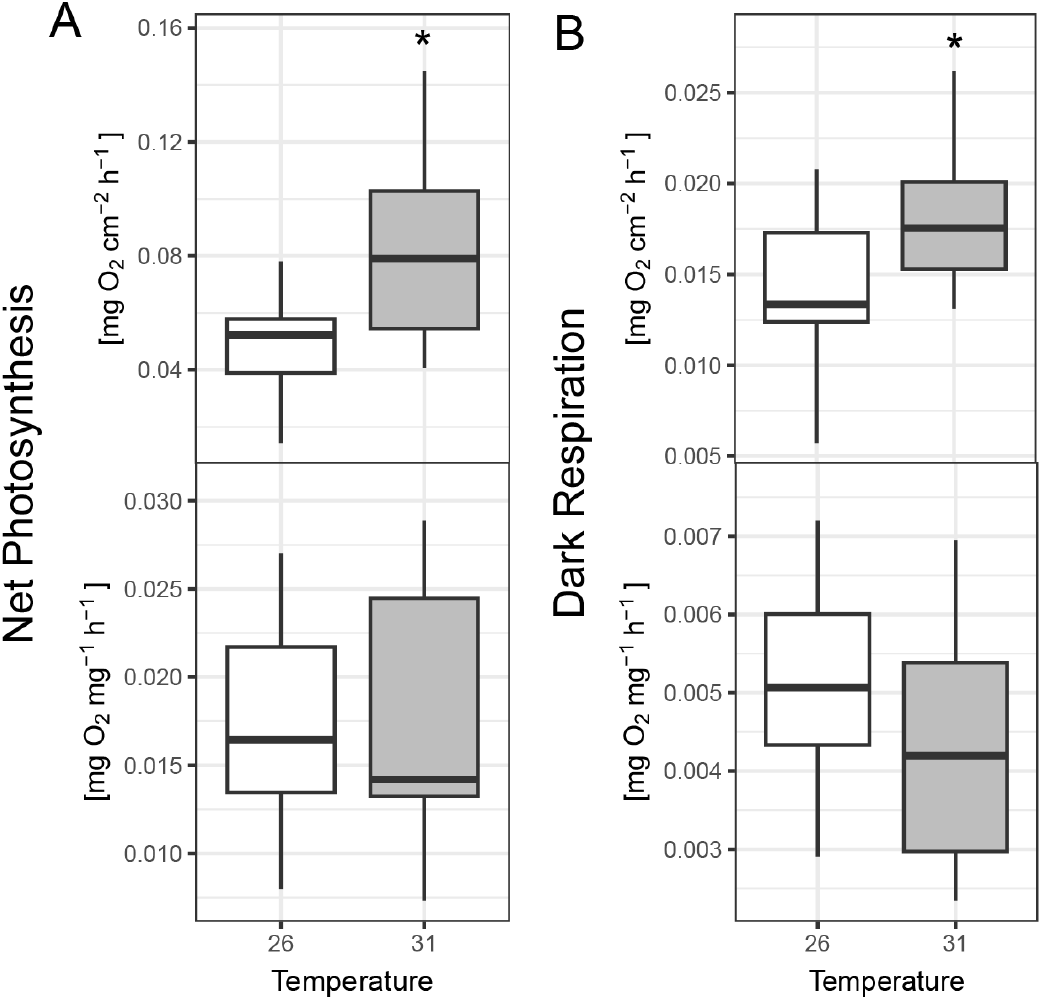
Metabolic performance of thermally acclimated corals. Live metabolic performance at holobiont level is shown as (A) net photosynthesis measured under light conditions and as (B) dark respiration. Both metrics are shown per cm^2^ of coral surface area (upper plot) and per mg of biomass weight (bottom plot). Thermal treatment in gray. Asterisks indicate significant group differences at significant levels: *p* < 0.05 (*). n = 11 to 12 per group.

### Skeletal growth and biomass accretion

Corals living at 31 °C calcified at a significantly slower pace (1.8-fold lower calcification rate) compared to corals living at 26 °C (Figure 2 A). At the same time, these corals formed skeletons of a higher densities at 31 °C (2.12 g cm^-3^) exceeding the densities measured in the ambient temperature group (1.58 g cm^-3^) by 1.4-fold (Figure 2 B). Living under the elevated temperature also resulted in a 2.5-fold significantly lower extension rate compared to corals living under the cooler ambient temperature (Figure 2 C). Overall biomass was significantly elevated in the coral holobionts at 31 °C (Figure 2 D, *p* < 0.01) as a result of significantly increased biomass in host (1.9-fold) and symbiont (1.5-fold) (Figure 2 E-F, *p* = 0.006, *p* = 0.009, respectively).

**Figure 2.**
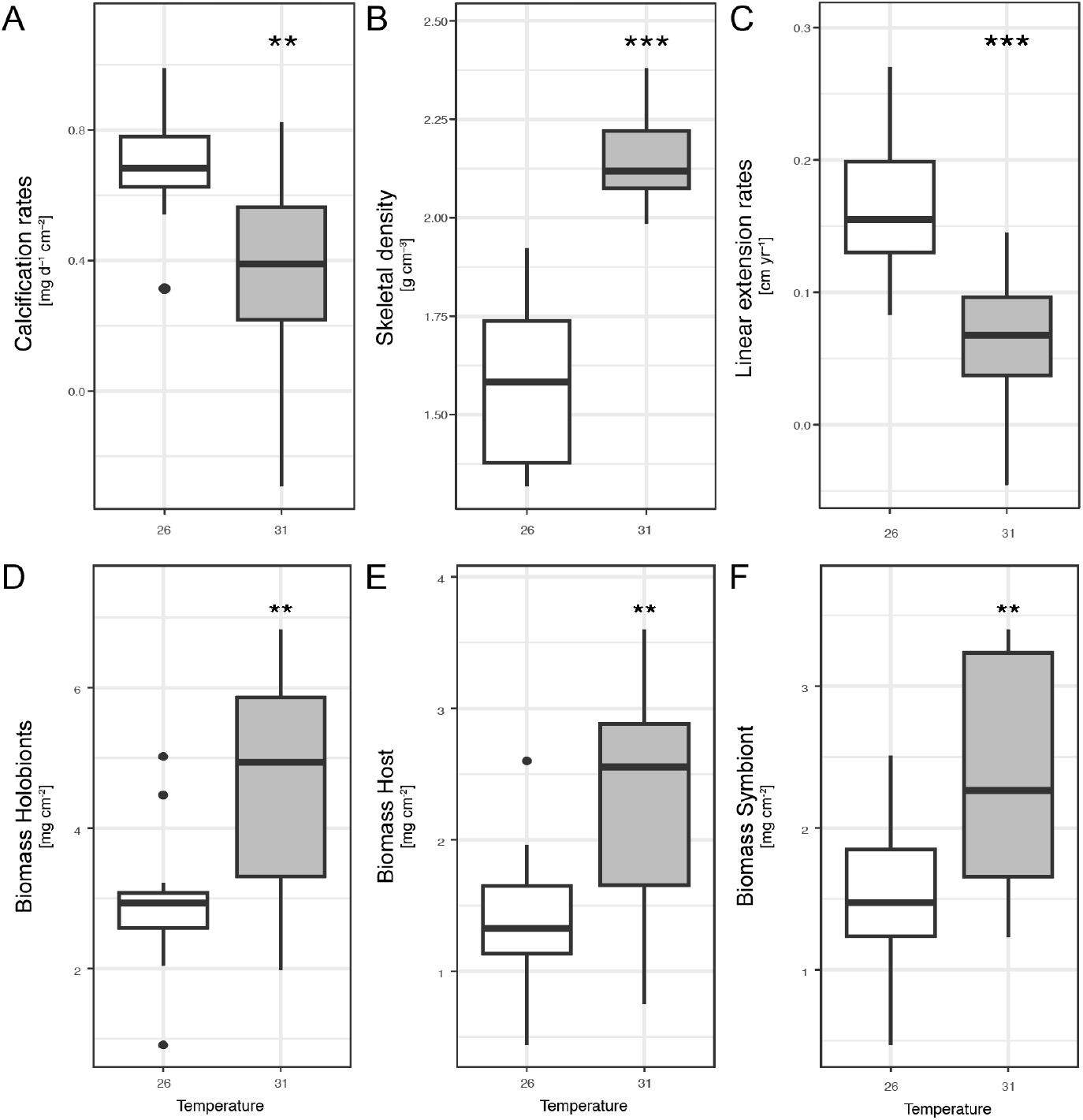
Growth traits of thermally acclimated corals. Skeletal growth parameters are presented as (A) calcification rates determined by buoyant weight measurements (as skeletal mass accretion per day and surface area), (B) skeletal density and (C) linear extension rates indicate the long-term growth trends of coral skeleton features. Biomass accumulation is shown (D) in total for the coral holobiont, and also specifically for the (E) host and (F) the symbionts. Extension rates were calculated assuming similar calcification rates along the entire surface area of individual fragments. Thermal treatment: Grey. Asterisks indicate significant differences at significant levels: *p* < 0.05 (*), *p* < 0.01 (**) and *p* < 0.001 (***); n = 11 to 12 per group.

### Energy storage

The total energy content of the holobiont was significantly higher in corals under the elevated temperature (1.8-fold increase, *p* < 0.001) when normalized to coral surface area (Figure 3 A). This difference was driven by the significantly increased lipid content in these corals (2.6-fold increase, *p* < 0.001 and *p* < 0.001, Figure 3 B). Carbohydrates and proteins remained at similar levels in both treatments (Figure 3 C-D). Proteins were slightly depleted in corals at 31 °C compared to corals at 26 °C (0.9-fold decrease, *p* < 0.05, when normalized to mg, Figure 3 D). The strongly increased lipid level determined at holobiont level at 31 °C was stemming from the host, which overall had significantly higher total energy content per surface area (median 59.0 J cm^-2^) under the elevated temperature (2.0-fold elevated, *p* < 0.001, Figure 3 E) and significantly higher lipid content (median 45.7 J cm^-2^ and 3.0-fold increase compared to 26 °C, *p* < 0.001, Figure 3 F). Furthermore, carbohydrate content per surface area were higher in host tissues under 31 °C (median 1.35 J cm^-2^, 1.2-fold elevated, *p* < 0.05, Figure 3 G), but protein content was at similar levels in both treatments (median 11.1 J cm^-2^ *vs.* 10.7 J cm^-2^, *n.s.*, Figure 3 H). Symbiont energy content played a proportionally smaller role in the total holobiont energy budget with values ∼10 - 25 J cm^-2^ and ∼5 - 13 J mg^-1^ (vs. host values ranging at ∼20 - 80 J cm^-2^ and ∼15 - 45 J mg^-1^). In comparison to the host and holobiont, symbiont energy content overall varied at a smaller scale between the thermal conditions of 26 °C and 31 °C, but the total energy content per biomass unit was significantly decreased under 31 °C (0.7-fold decrease, *p* < 0.01, when normalized to mg of biomass, Figure 3 I), which is in contrast to what we have found at the holobiont and host level. This decline was driven by significant declines per unit of biomass revealed for all three parameters, lipids (0.7-fold, *p* < 0.05, Figure 3 J), carbohydrates (0.7-fold, *p* < 0.05, Figure 3 K), and proteins (0.5-fold, *p* < 0.01, Figure L). When calculated by surface area all the three symbiont parameters including symbiont total energy reserves appear homogeneous between the two thermal conditions (total median 16.2 J cm^-2^ at 31 °C, 15.3 J cm^-2^ at 26 °C, *n.s.*).

**Figure 3.**
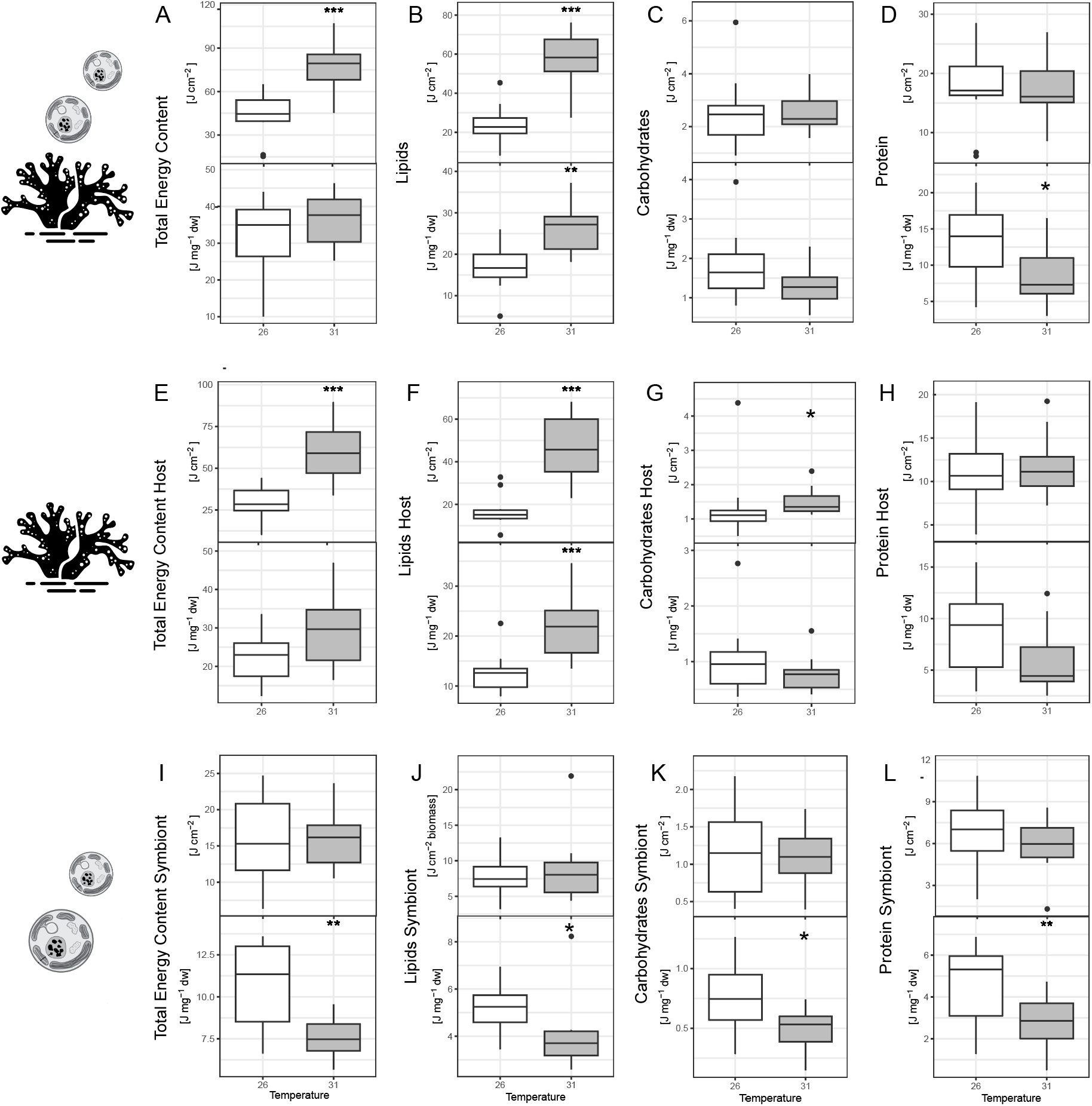
Tissue energy content of thermally acclimated corals. The total energy reserves and content of lipids, carbohydrates and proteins in coral tissues are shown for (A - D) the holobiont and also for (E - H) hosts and (I - L) symbionts, individually. All variables are shown per cm^2^ of coral surface area (upper plot) and per mg of biomass weight (bottom plot). Thermal treatment in gray. Asterisks indicate significant group differences at significant levels: *p* < 0.05 (*). n = 11 to 12 per group.

## Discussion

Our study reports first insights into the metabolic shifts and trade-offs in pocilloporid corals that acclimated to elevated temperatures relative to their reef of origin and have remained under these conditions for over six years. We observed that corals maintained at the cooler ambient (26 °C) and the elevated (31 °C) temperature for six years appeared visually healthy and thriving. The 31 °C-acclimated corals operated at increased metabolic rates, while prioritizing energy investment into lipid storage and biomass accumulation over skeletal growth. These acclimated corals hosted symbionts that appeared compromised (i.e., lower energy content) in comparison to the corals kept at 26 °C. We discuss the observed differences and trade-offs and their consequences for this globally abundant and ecologically relevant coral species under an elevated temperature, including the potential long-term consequences for the reef ecosystem.

### Shift of energetic production and investment under elevated temperature

Marine invertebrates can maximize their fitness under challenging environmental circumstances through prioritizing one trait over another. They undergo physiological shifts that change their relative energy allocation strategy (Zera & Harshman, 2001). Our data highlight that long-term exposure of corals to an elevated temperature can result in a remarkably strong channeling of resources into tissue growth and accumulation of energy reserves, while neglecting growth of coral colonies. We must assume that energy expenditures for biomass were significantly increased under the elevated temperature at the expense of skeletal growth. Since tissue growth is more energy consuming than skeletal accretion (Kenneth R. N. Anthony et al., 2002), corals with enhanced productivity at 31 °C were likely able to cover the increased costs for biomass, but this might have led to a deficit in meeting energy requirements for calcification. In other previous studies, however, under various challenging environmental conditions other than elevated temperature, skeletal growth was typically prioritized. In particular under light deprivation which slows photosynthetic rates, the coral *Montipora digitata* shifted its relative energy investment from tissue growth to reproduction and skeletal growth as a response to declining resource availability (Leuzinger et al., 2012). Similarly, under severe energy limitation caused by shading, this coral maintained skeletal growth even at the expense of reproduction. In other cases, biomass accumulation was reported to increase under more light availability in *P. acuta*, while calcification rates remained stable (C. B. Wall et al., 2017). However, across a diversity of coral species including Pocilloporidae, tissue biomass has been typically negatively correlated with skeletal growth and most often, the slow-growing coral species would maintain more biomass per surface area (Precoda et al., 2020). This aligns with the observation in our corals, where *P. acuta* shifted from fast colony growth to slow growing under elevated temperature, at the same time strongly increasing their biomass, as a possible acclimation strategy. In conclusion, the pattern appears that maintaining skeletal growth under resource constraints (e.g., light- or nutrient-deficient conditions) is preferred over biomass accumulation. On the other hand, a productivity boost (e.g., increased photosynthesis and/or algal symbiont density) associated with increasing temperatures tends to promote the investment into biomass and lipid accumulation at the expense of skeletal growth (Anthony et al., 2002; Tanaka et al., 2007), as was the case in this study.

### Benefits of the investment shift into biomass and energy reserves

We observed that biomass composition of corals differed between the two thermal regimes. The 2.6-fold increase of the tissue lipid fraction in corals living at the elevated temperature shows that they prioritized energy investment into lipid storage. This acclimation trait observed in our 6-year long study does not align with observations of short-term studies, where elevated temperatures caused a depletion of tissue lipids in corals (e.g., Bove et al., 2022; Schoepf et al., 2013). In these short-term bleaching experiments, depletion of host tissue lipids should be interpreted as a stress-response driven by a shift in symbiont cellular pathways (gluconeogenesis, i.e., glucose production via lipid and amino acid breakdown), and consequent change in the quality of translocated products (i.e., decrease in fatty acids and complex molecules) (Hillyer et al., 2017; Pei et al., 2022). This highlights the importance of studying acclimation of organisms over their relevant biological scales, where successful acclimation mechanisms, which may include trade-offs, can be distinguished from stress-responses.

By increasing their tissue energy content compared to cooler temperature controls, corals in our experiment have likely gained the benefit of preparedness for future unfavorable conditions, as high lipid stores have been previously linked to better coral health, lower mortality and higher recovery rates following stressful conditions (Anthony et al., 2002, 2009). For example, lipid compounds are utilized first during the onset of bleaching (Grottoli et al., 2004; Rodrigues et al., 2008) and, thus, a high energy content can enable corals to withstand bleaching conditions for a longer period of time. Furthermore, investments into tissue accumulation and energy content can be beneficial by enabling rapid tissue repair after events of stress and tissue damage (Henry & Hart, 2005; Traylor-Knowles, 2016). It is a common notion that rising environmental temperatures accelerate biochemical and metabolic reactions in marine ectotherms (Angilletta et al., 2004; Corkrey et al., 2014), which in corals is often accompanied by increasing investment into cell protection and tissue maintenance (to avoid cell damage), while colony growth is reduced (Hornstein et al., 2018). Previous studies have shown enhanced investment into higher antioxidant activity and increased biomass content in *Montipora capitata* after repeated thermal stress (C. B. Wall et al., 2018, 2021). Such progressive upregulation of constitutive antioxidant activity (e.g., superoxide dismutase and catalase content) typically helps to protect tissue biomass (Lesser et al., 1990) which can increase the potential for overall survival under thermal stress.

The question of whether large energy reserves expectedly accompanied by antioxidant frontloading are fundamentally beneficial to corals under extreme events such as bleaching remains unanswered. Despite energy reserves positively correlated with bleaching resistance and recovery capacity (Anthony et al., 2009; Grottoli et al., 2014; Hoegh-Guldberg, 1999), in some cases, bleaching resistance has been found to be decoupled from the levels of energy reserves in corals (Grottoli et al., 2004; Precoda et al., 2020). In the present study, we did not examine whether energy reserves in warm-acclimatized *P. acuta* individuals would be beneficial during acute heat stress. This should be an important next step in the study of these acclimated corals together with the assessment of their antioxidant activity.

### Reduced skeletal growth and consequences

Considering that energy supply is typically sufficiently high to cover all physiologically relevant processes in marine ectotherms under the moderate thermal conditions (Leuzinger et al., 2012; Sokolova et al., 2012), the reduction in calcification at the elevated temperature in this experiment is an indicator that corals were operating beyond their thermal optimum for skeletal growth, where energetic trade-offs occur. Despite their visually healthy appearance and remarkable performance regarding biomass accumulation, the temperature of 31 °C seems to pose a challenge, potentially presenting a suboptimal thermal environment for these corals. The observed response of skeletal growth was in agreement with the thermal optimum ranging between 27.5 - 29.5 °C that is known for a range of coral taxa from the Great Barrier Reef (GBR) or the Caribbean (Álvarez-Noriega et al., 2023; Silbiger et al., 2019). For *P. verrucosa* from the GBR, for instance, optimal calcification temperature was 29.5 °C and severe declines in calcification capacity have been noted beyond this optimum, with up to ∼30 % declines already under 31 °C (Álvarez-Noriega et al., 2023). A similar situation can be assumed for the corals in the present study, where calcification rates at 31 °C were 40 % lower compared to the ambient temperature conditions. Since our corals’ home reef, Luminao, is a fringing reef that can experience midday temperature peaks above ∼31 °C during the hottest months of the year, the new constant exposure temperature of 31 °C in our experiment was expected to exceed their natural thermal optimum. In a recent study, exposure of *P. damicornis* corals to 31 °C clearly exceeded the growth optimum as indicated by the reduced growth rates which was accompanied with the impairment to control their calcifying fluid (Guillermic et al., 2021). Such inability to maintain control over the calcifying fluid condition, may also account for the observed lowered calcification rates in our study under the elevated temperature.

Furthermore, our findings revealed a temperature-induced change in skeletal properties. Corals at 31 °C had carbonate skeletons of higher density, which should provide them with a higher skeletal robustness. Most commonly the slow-growing coral species would form higher density skeletons (Precoda et al., 2020). Within one coral species the tendency for higher-density skeletons is known for colonies that inhabit challenging environments like high energy habitats such as the reef crest, where physical forcing through wave and current impact is high (Madin et al., 2008; Scoffin et al., 1992; Smith et al., 2007). This is undoubtedly beneficial in environments under physical forcing, unlike our coral aquaria. Interestingly, corals in thermally challenging environments, such as inshore reefs, are more commonly known for their reduced skeletal density (McWilliam et al., 2022). The only exception in that multi-species study was *P. cf damicornis*, supporting our finding of higher density skeletons under elevated thermal conditions and also demonstrating that such growth tendencies, and consequently trade-offs, are likely species-specific. Corals with high-density skeletons must calcify faster in order to keep up with the skeletal linear extensions achieved by corals with lower-density skeletons. Hence, the investment into a dense skeleton comes at a cost of reduced linear extension at a similar growth rate, resulting in slower colony expansion (Precoda et al., 2020). In our study, high density multiplied by the lower calcification rates of corals at 31 °C, resulted in skeletal extension rates substantially lower compared to their 26 °C counterparts. In this context developing high density skeletons, especially in combination with lower calcification rates, needs to be considered as a significant trade-off with ramifications not only at holobiont scale, but also far-reaching ecological consequences for reef growth dynamics and maintenance of the three-dimensional reef structure.

### Changes in host-symbiont relationship

Unlike calcification rates, the photosynthetic performance of symbionts was not constraint under the elevated temperature. In contrast, photosynthesis was boosted in the 31 °C-acclimated coral in our study. This aligns with the results from short-term coral performance assays conducted in the Caribbean (Silbiger et al., 2019) and underscores that optima for photosynthetic productivity are not constrained at the elevated temperatures tested. However, the photosynthetic boost, which is suspected to increase overall energy levels for the holobiont, was not accompanied by any increase of symbiont biomass nor any change of their energy content in our study. Instead, only the host tissues were able to increase biomass remarkably, suggesting that the transfer of energy from symbiont to host under elevated temperatures must have been increased, either by optimizing or enforcing translocation of photosynthates. It has been previously established in a pocilloporid coral that a ‘sub-lethal/sub-bleaching’ thermal exposure had a significant impact on nutrient cycling and metabolism, entailing modifications of the energetic exchange of the two partners in symbiosis (Gibbin et al., 2018). Additionally it has been shown that increased photosynthesis can be coupled with a significant increase in heterotrophic feeding rates in a cnidarian holobiont (Leal et al., 2015), presenting another possible contribution that might have further fueled growing host energy storage in our experiment.

The detailed examination of coral tissues by host and symbiont fraction, allowed to further obtain a glimpse into the complex dynamics of possible symbiont-host interactions that accompanied the thermal acclimation. Symbionts at 31 °C were diagnosed with lower protein, carbohydrate, and lipid levels per symbiont biomass. Interestingly, these values, in relation to coral surface area, have remained similar under both temperatures. This shows that symbiont biomass per host biomass did not change, despite boosted energy production and once again highlights the strong investment and resource channeling into the energy storage of the host. These nuanced findings further indicate that symbionts likely underwent cell-morphological changes influenced by temperature. The capacity of morphological restructuring has been reported from symbionts that were classified as stress “resistant” compared to other more “sensitive” species/strains, which did not feature such morphological plasticity (Hoadley et al., 2015). Resistant symbionts were not only able to increase their own protein and lipid storage, but also demonstrated morphogenesis (enlargement) of chloroplasts at elevated temperature, as well as an increase in cell volume, chlorophyll fluorescence, and pigment content (Gong et al., 2020; Hoadley et al., 2016). As such, these symbionts may have increased their chloroplast volume to increase and optimize their photosynthetic output under the elevated thermal conditions that contributed to boosting the metabolic rates in both holobiont partners. Their plasticity coupled with increased energy content, observed in these previous studies, can be interpreted as a beneficial trait of the symbionts, which can help enhance holobiont stress resistance under challenging thermal conditions. In contrast, our findings show a ‘skinny’, but productive symbiont paired with a well-nourished host, highly enriched in tissue lipids, which also can be an indication of a changed nutrient cycling between two partners (Gibbin et al., 2018) and of an enhanced translocation of symbiont resources (Rädecker et al., 2021). Despite the fact that different symbiont species (or strains) maintain distinct metabolic traits and can employ different energy/nutrient transfer strategies (Leal et al., 2015), the here recorded differences in symbiont properties, likely do not reflect different symbiont species. At the age of one year, all corals used in our experiment harbored the same dominant symbiont species, *S. durusdinium* ‘D1/D2d’, with no differences in symbiont assemblages between the temperature treatments (unpublished data). Parent colonies in Guam hosted the same D1/D2d strain and throughout the first 12 months, no changes in symbiont assemblages were detectable. While most studies to date have investigated the transition period between the stable and the unstable symbiotic state during thermal stress (aka. coral bleaching), our study provides new valuable insights into the symbiont-host trait dynamics in a stable symbiosis that has acclimated to an elevated temperature of 31 °C. We do not fully understand yet, whether this 31 °C-acclimated symbiotic state will also prove beneficial during an acute thermal stress event. We can hypothesize two contrasting scenarios, 1) that the increased investment into host tissues will increase stress resistance and will help the coral to deal with future stressors (Grottoli et al., 2004), or, 2) that the enhanced translocation of symbiont resources to the host brings the holobiont closer to a dysbiotic state (Rädecker et al., 2021) and, thus, will increase its susceptibility to stressors. This remains to be determined in a future study, but overall, our current findings have already shed light in the physiological and metabolic shifts that allow coral holobionts to acclimate successfully under warmer temperatures.

### Underlying mechanisms of observed coral responses under elevated temperature

In this study we describe the successful acclimation of *P. acuta* to the new conditions of an elevated temperature, which could be a result of physiological plasticity, genetic selection and adaptation, or a combination of both (Chevin & Hoffmann, 2017; R. J. Fox et al., 2019; Kelly, 2019; Palumbi et al., 2014; Torda et al., 2017). We suspect that the thermal history of the parental colonies in the field, as well as the early exposure to elevated temperatures of the offspring, have contributed to the acclimation success of corals in our experiment. Since exposure to thermal variability is a good predictor of high stress-resistance and large plasticity in corals (Hackerott et al., 2021; Rivest et al., 2017; M. Wall et al., 2023), the thermal history of the parents from the Luminao reef flat, which has a large thermal range, could be one explanation, why the offspring was able to acclimate to the new elevated temperature of 31 °C. Furthermore, corals in this study have “learned” to thrive under the new elevated temperature, since the very first exposure at a juvenile stage, as no signs of distress were noted during the six years of cultivation. This early exposure to the elevated temperature during their recruitment might have promoted the success of acclimation, as developmental exposure to certain drivers like an elevated temperature have been shown to influence plasticity in various organisms (Bowler & Terblanche, 2008). However, it will be worthwhile to further explore the underlying genetic make-up of the offspring by investigating whether differences in allele frequency can be identified between the two groups, since allele shifts were often associated with enhanced thermal tolerance of *ex situ* bred corals (Dixon et al., 2015; Howells et al., 2021; Quigley & van Oppen, 2022). In our experiment, the possibility remains that selection of recruits took place right after settlement, since a higher number of recruits survived under 29 °C compared to 31 °C (Supplementary Figure S2). Evolutionary processes cannot yet be fully ruled out as a driver for the observed physiological differences reported between the corals raised at the two thermal regimes. Larval selection process has been characterized in other studies showing that heat-selected coral larvae were significantly enriched in heat-shock proteins and had improved energy production and conversion, as well better oxidative stress and immune responses (Dixon et al., 2015; Howells et al., 2021). Hence, to fully elucidate to what extent our observations were mainly due to physiological plasticity or associated with the genotypic composition of the coral offspring, additional genotype analyses of offspring and ideally of the source population will be required.

### Ecological implications and considerations for active reef restoration

The increasing severity and frequency of deteriorating coral bleaching events (Donner et al., 2005; van Hooidonk et al., 2013) have been driving the development of proactive measures that aim to protect corals from thermal stress (van Oppen et al., 2015). Some of the anticipated approaches consider selection of thermally tolerant coral specimens for reef restoration (Humanes et al., 2021; Morikawa & Palumbi, 2019), while others intend to use thermal preconditioning treatments aiming to improve thermal tolerance of nursery corals (DeMerlis et al., 2022; Hancock et al., 2021; Henley et al., 2022; M. Wall et al., 2023). However, our findings have demonstrated that the desired trait of higher thermal tolerance can come at the cost of skeletal growth, specifically for the coral *P. acuta* (from Guam). Further trade-offs beyond the decline of colony growth are possible. It will be crucial to investigate reproduction, as it determines coral population fitness with critical repercussions for the persistence of reef communities and the recovery of populations following severe heat stress events (Fisch et al., 2019; Johnston et al., 2020; Levitan et al., 2014).

The far-reaching ecological consequences of trade-offs have not been considered nor assessed yet. For instance, a reduction in skeletal growth is expected to limit the growth capacity of a whole reef structure, which is a critical ecological feature ensuring that a reef will be able to maintain a positive carbonate budget (Roik et al., 2015, 2018), keep up with future sea-level rise (Perry et al., 2018), and hence offer coastal protection and retain its ecosystem services and in the future (Eddy et al., 2021; IPBES, 2019). Furthermore, with reduced colony growth rates corals may show less resilience and poor potential of recovery from the pressures of other stressors, such as, i.e., increased forcing and frequency of storms and ocean acidification which increases with ocean warming (Madin et al., 2014; McCulloch et al., 2012). On the other hand, corals that are able to produce a skeleton of higher density, such as observed in the heat-acclimated *P. acuta* in this study, may be able to buffer some of the negative effects to ocean acidification, which has been demonstrated to reduce coral skeletal density (Mollica et al., 2018). Consequences of trade-offs can be complex and whether adaptation/acclimation to one stressor (e.g., temperature) may also increase the resistance to other stressors (e.g., ocean acidification, eutrophication, disease), is a question that has so far received little attention. Trade-offs could, both, hamper or improve the success of current interventions and reef restoration efforts that desire to increase the thermal tolerance of corals. Some restoration projects have already integrated an assessment of trade-offs in their monitoring programs, e.g., coral nurseries in Florida reported a potential trade-off between disease resistance (a desired trait) and reproductive output of their nursery corals (Koch et al., 2022). Overall, studies exploring trade-offs of coral thermal tolerance underscore the importance of taking a holistic view on this matter. Currently, they show that an efficient strategy to create new intervention protocols should focus on a set of multiple desired traits for coral restoration recruits (Caruso et al., 2021; Edmunds & Putnam, 2020; Wright et al., 2015). Wright et al. (2019) research provided the first indication that certain coral traits could be advantageous against multiple stressors, however it is noteworthy that the traits underpinning stressor tolerance were not identified and the experiment only lasted 10-days. Careful consideration, assessment, and cost-benefit evaluation of each new method and of the full suite of potential ecological consequences, which may arise from the method, will be vital to the development of efficient new interventions.

## Conclusion

Knowing that reef-building corals have the capacity to acclimate to elevated temperatures, we have set out to determine if such increases in thermal resistance come at any costs for a coral. Examinations of physiological and metabolic features of corals acclimated under two distinct thermal conditions (a cooler ambient and an elevated temperature) have identified two key trade-offs. After six years at the elevated temperature, corals allocated more resources towards soft tissue growth and lipid storage, while maintaining slower yet denser skeleton growth. The trade-off between energy storage and skeletal growth, likely involved the exploitation of symbionts, demonstrating how corals need to balance physiological and metabolic mechanisms in order to acclimate to higher temperatures. On the one hand, the coral hosts at the elevated temperature appeared well prepared to withstand future stressors thanks to their energy reserves. On the other hand, their symbionts were unable to accumulate substantial energy stores, potentially rendering them more vulnerable. Our results demonstrate how a “gain” in thermal tolerance could hinder the calcification and reef-building potential of corals, while enhancing the coral host’s resilience to stressors. Further long-term assessments of trade-offs in other coral species are needed to determine if these trade-offs are specific to *P. acuta* or more widespread. Our results challenge the observations of short-term studies, where elevated temperatures depleted tissue lipids in corals, emphasizing the significance of studying acclimation over relevant biological scales. Long-term studies like ours will help to obtain a more comprehensive picture of the future coral reef trajectory and help to more accurately assess the potential of anticipated interventions that aim at increasing coral thermal tolerance.

## Supporting information

Data_sharing_Tables

## Funding and acknowledgments

AR and JS acknowledge the funding of the Helmholtz Institute for Functional Marine Biodiversity at the University of Oldenburg, Niedersachsen, Germany. HIFMB is a collaboration between the Alfred-Wegener-Institute, Helmholtz-Center for Polar and Marine Research, and the Carl-von-Ossietzky University Oldenburg, initially funded by the Ministry for Science and Culture of Lower Saxony and the Volkswagen Foundation through the “Niedersächsisches Vorab” grant program (grant number ZN3285). PJS acknowledges support via startup funding by the Institute of Chemistry and Biology of the Marine Environment (ICBM, University of Oldenburg). Funding for the initial research that produced the corals used in this study was provided by the German Academic Scholarship Foundation. We thank M.Sc. Tabea Platz for her assistance in the wet laboratory during physiological experiments and Esther Lüdtke as well as Irini Kioupidi for support in coral tissue processing. Thanks to Prof. Gabriele Gerlach and Susanne Wallenstein for providing laboratory space and equipment to perform live physiology and skeletal density measurements. SN thanks Dr. Mareen Möller for her support in producing and rearing the corals used in this study. We thank Dr. Laurie J. Raymundo (University of Guam Marine Laboratory) for providing temperature data from the “home” reef of our corals.

## Permissions

Research was conducted under the permit of the DEPARTMENT OF AGRICULTURE DIVISION OF AQUATIC AND WILDLIFE RESOURCES (DAWR) and MPA APPLICATION SPECIAL REQUEST (Section 63123 of Title 5, Guam Code Annotated GCA) to PJS. Corals were collected under the Special License For The Collection Of Coral, issued to UOGML by DAWR under section 63123 of Title 5, GCA, and exported under permission of CITES issued by the US Fish and Wildlife Service (export # 15US62023B/9).

## Authors’ contributions

AR and JS conceived the study, designed and performed the experiments. SN collected, reared and provided corals used in this study. SN and MJ maintained aquaria facilities during experimental work. JS, AR, MD, MR, DB, AF, MW performed experiments and laboratory work. PJS provided facilities, funding, and laboratory space. MW, AR, MD, AF analyzed and visualized data. AR, MW, JS wrote and edited the manuscript. MR, SN, AF, PJS read and edited the manuscript. All authors read and approved the final manuscript.

## Availability of data and materials

All data is included in the supplementary material to this paper.

## Declarations

The authors declare that they have no competing interests.

## Notes

### Competing Interest Statement

The authors have declared no competing interest.

### Summary of Updates

We updated the Supplementary data sheets by collating all relevant original and analysis data sheets in one file.

