## Supplementary material for "Trade-offs in a reef-building coral after six years of thermal acclimation": Supplmentary_Materials_PDF

#### Supplementary Materials

#### *Text*

##### Text S1 Phylogenetic analysis of corals

DNA was extracted from coral tissue samples using a Macherey-Nagel NucleoSpin Plant II Kit (Macherey-Nagel GmbH & Co. KG, Düren, Germany). A mitochondrial open reading frame (mORF) was amplified using primers developed and described by (Flot & Tillier, 2007) for corals of the genus *Pocillopora* (FATP6.1 5′-TTTGGGSATTCGTTTAGCAG-3′ and RORF 5′-SCCAATATGTTAAACASCATGTCA-3′). PCR reactions (30 µl) contained 15 µl Qiagen Multiplex PCR Kit (QIAGEN®), 1.88µl primers (each primer at 20µM, final concentration 1.25µM), 8.74 µl NFW and 2.5 µl template DNA (10 – 40 ng/µl). Cycle conditions were: 95 °C for 15 min + 35 x [94 °C for 60 s, 53 °C for 60 s, 72 °C for 60 s] + 72 °C for 10 min. 25 µl of each PCR product was sent to the Institut für Klinische Molekularbiologie (IKMB) at the Christian-Albrechts-Universität zu Kiel for PCR clean up and bi-directional Sanger sequencing (ABI 3730xl DNA Analyzer). The respective forward and reverse sequences were assembled, aligned, and trimmed in CodonCode Aligner v9.0.1.3 (codoncode corporation). The contigs were checked for gaps and insertion/deletion polymorphisms (indels). Results were transferred into BLAST® and all alignments of unambiguous, high quality reads were converted in MEGA 11 (Tamura et al., 2021) for further analysis of distinct haplotypes in DNAsp. Thereafter, the resulting NEXUS file was exported into PopART (Leigh & Bryant, 2015). For the median-joining haplotype network, sequences of the sister species *P. verrucosa/P. meandrina* from the Indo-Pacific and *P. damicornis* from the Ningaloo Reef were obtained from GenBank (Pinzón et al., 2013; Thomas et al., 2014).

Unambiguous, high-quality alignments (821 bp fragments) of a mitochondrial open reading frame (mORF; accession numbers: OP776415 - OP776438) were identified as *Pocillopora acuta* and compared to GenBank sequences of *P. damicornis type β,* now *P. acuta* (accession numbers: KT879932 - KT880039) and *P. acuta* (accession numbers: MH064356 - MH064390). The data set provided one distinct haplotype (Figure S3).

### *Figures*

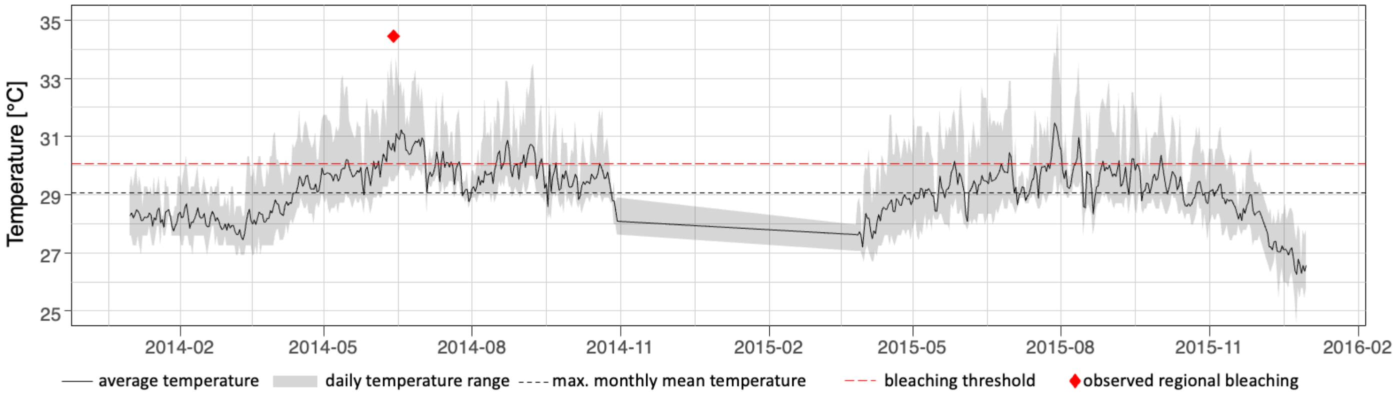


**Figure S1 Reef temperature recorded in 2014-2015 at the coral collection site.** Average reef temperature and diurnal temperature range (gray area illustrating the min to max recorded temperature) recorded at the Luminao reef flat, Guam in 2014 to 2015. Dashed lines represent the maximum monthly mean temperature (black) and bleaching threshold (red) for the region (Data by Raymundo Lab, (University of Guam, Marine Laboratory). Observed regional bleaching is indicated by red diamond.

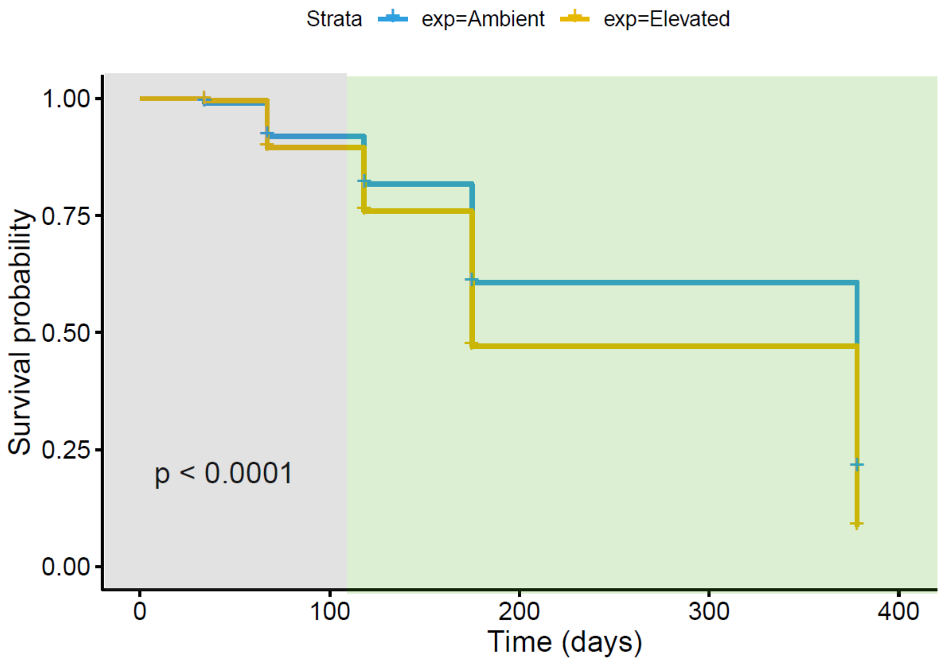


**Figure S2 Coral recruit survival after settlement of coral larvae in Guam in 2015.** Survival of coral recruits was tracked for over one year after settlement, starting in July 2015 and ending in August 2016 (i.e., before the start of the experiment in 2021). The Kaplan-Meier Survival plot visualizes the survival probability of individual corals at a specific time point over the monitoring period. Grey shading indicates the time span, when corals were maintained in a flow-through system in Guam. Green shading indicates the time span, when corals were maintained in the facility in Wilhelmshaven, Germany. Time on the x-axis is provided in Days. “Ambient” = Ambient temperature conditions of 29 °C; “Elevated” = Elevated Temperature conditions of 30 - 31 °C.

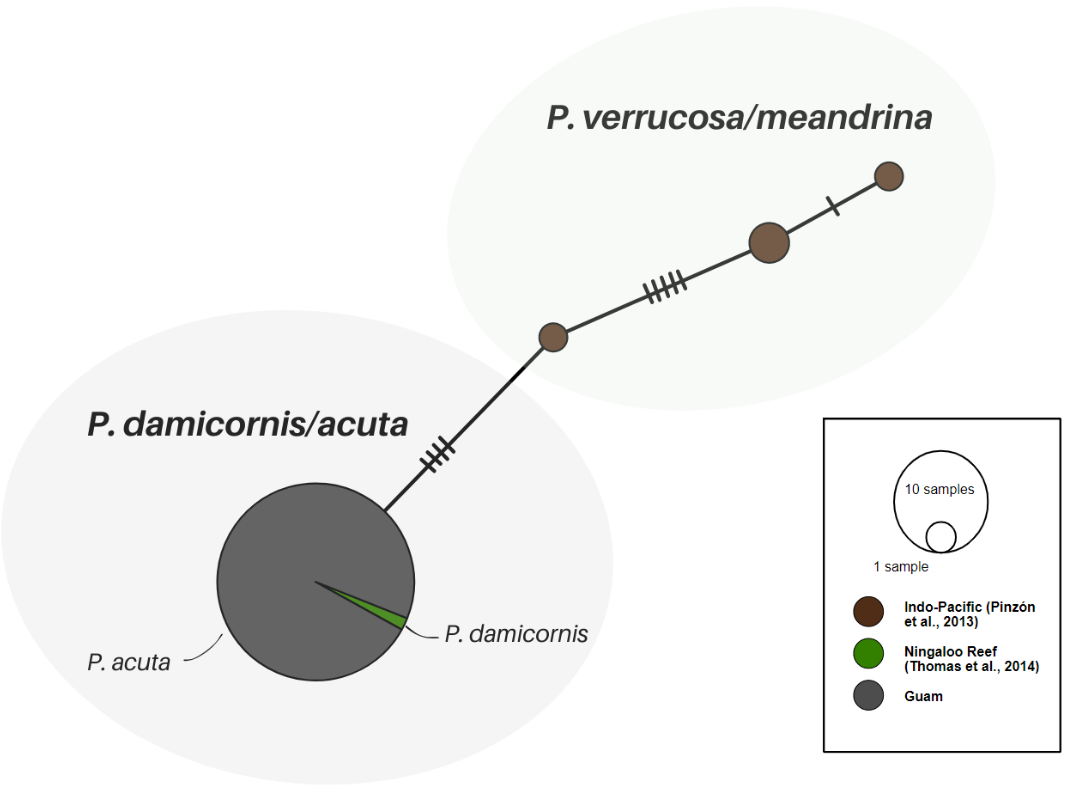


**Figure S3. Phylogenetic analysis of pocilloporid corals from Guam.** Minimum-spanning haplotype network of mORF DNA sequences from Guam (gray circle). The alignment consisted of 24 sequences, 835 bp length. Sequences were referenced against one distinct *P. damicornis* haplotype found on the Ningaloo Reef (green coloring; Thomas et al., 2014). *P. verrucosa/meandrina* references are from the Indo-Pacific (brown circles) identified by Pinzón et al. (2013). */**Sequences obtained from *GenBank*.
